# A cross-species assessment of behavioral flexibility in compulsive disorders

**DOI:** 10.1101/542100

**Authors:** Nabil Benzina, Karim N’Diaye, Antoine Pelissolo, Luc Mallet, Eric Burguière

**Affiliations:** Institut du Cerveau et de la Moelle épinière, ICM, Inserm U 1127, CNRS UMR 7225, Sorbonne Université, F-75013, Paris, France; Assistance Publique-Hôpitaux de Paris, DMU IMPACT, Département Médical-Universitaire de Psychiatrie et d’Addictologie, Hôpitaux Universitaires Henri Mondor - Albert Chenevier, Université Paris-Est Créteil, Créteil, France; INSERM U955, IMRB, Créteil, France; Department of Mental Health and Psychiatry, Global Health Institute, University of Geneva, Geneva, Switzerland

**Keywords:** Behavioral flexibility, Compulsions, OCD, *Sapap3* KO, translational research

## Abstract

**Background:** Compulsive behaviors, one of the core symptoms of Obsessive-Compulsive Disorder (OCD), are defined as repetitive behaviors performed through rigid rituals. The lack of behavioral flexibility has been as being one of the primary causes of compulsions, but studies exploring this dimension have shown inconsistencies in different tasks performed in human and animal models of compulsive behavior. The aim of this study was so to assess the involvement of behavioral flexibility in compulsion, with a similar approach across different species sharing a common symptom of compulsivity.

**Methods:** 40 OCD patients, 40 healthy individually matched control subjects, 26 C57BL/6J *Sapap3* KO mice and 26 matched wildtype littermates were included in this study. A similar reversal learning task was designed to assess behavioral flexibility in parallel in these two species.

**Results:** When considered as homogeneous groups, OCD patients and KO mice expressing compulsive behaviors did not significantly differ from their controls regarding behavioral flexibility. When clinical subtypes were considered, only patients exhibiting checking compulsions were impaired with more trials needed to reach the reversal criterion. In KO mice, a similarly impaired subgroup was identified. For both species, this impairment did not result in a greater perseveration after reversal, but in a greater lability in their responses in the reversal condition. Moreover, this impairment did not correlate with the severity of compulsive behaviors.

**Conclusions:** In our cross-species study, we found no consistent link between compulsive behaviors and a lack of behavioral flexibility. However, we showed in both species that the compulsive group was heterogeneous in term of performance in our reversal learning task. Among the compulsive subjects, we identified a subgroup with impaired performance not due to perseverative and rigid behaviors as commonly hypothesized, but rather to an increase in response lability.

## INTRODUCTION

A full understanding of the underlying neural and cognitive bases of obsessive-compulsive disorder (OCD) remains a major challenge. The disorder affects 2-3% of the general population (1) and is characterized by obsessions (intrusive and unwanted thoughts) often associated with compulsions (repetitive and ritualized behaviors or mental acts). From a neurobiological perspective, OCD is characterized by a prefronto-striatal circuits dysfunction (2). However, the link between these dysfunctions and both the clinical expression of OCD and its genetic basis is poorly understood; especially since the latter remains unclear (3). It may therefore be more appropriate, and feasible, to look at other mammalian species which could share the neural and genetic bases of the disorder (4).

Behavioral flexibility is a cognitive function which can be studied in detail in other species. It refers to the ability to flexibly adjust behavior to a changing environment (5), with its impairment echoing the rigid compulsive behaviors in OCD patients who are resistant to change. However, there is no consensus that such a deficit exists in OCD with inconsistencies between studies (6) using reversal learning (7, 8), task switching (9, 10), or intra/extra-dimensional set shifting (11, 12) paradigms. Beyond methodological considerations like small sample sizes (13, 14), the clinical heterogeneity of OCD patients may have contributed to these discrepant results (15–19). In contrast, activation of OCD patients’ prefrontal regions has been more consistently reported as dysfunctional while performing flexibility tasks, in particular the orbitofrontal cortex (OFC) (20, 21). These prefrontal areas are parts of the limbic and associative cortico-basal ganglia circuits where abnormalities have been well characterized in OCD patients, either in the resting state or during symptom provocation (2, 22). Similar observations have been recently made in the *Sapap3* KO mice, the current predominant animal model of compulsive behavior (23). These mice have been genetically engineered and lack the SAP90/PSD95-associated protein 3 (*Sapap3*), a postsynaptic scaffolding protein mainly expressed in the striatum (24). This mutation results in expression of compulsive grooming behaviors underpinned by neurophysiological impairments of the prefronto-striatal circuits (23, 25), including the OFC (26). Additionally, their excessive grooming behaviors can be significantly reduced after chronic administration of fluoxetine as observed in OCD patients (23). These observations emphasize the relevance of the excessive grooming behaviors observed in *Sapap3* KO mice as analogous to compulsions observed in OCD patients. Regarding behavioral flexibility, two recent studies (27, 28) have studied these mice in a spatial reversal learning task. The results suggest some deficits, although of different nature in the two studies, but comparison is difficult since the experimental paradigm is different from that used in humans.

To improve the comparison of human and animal results, similar procedures need to be developed across species to ensure similar task parameters and psychometric properties (29, 30), and to reinforce the comparative value of the results (31). Concerning behavioral flexibility, assessments rely in most cases on spatial discrimination tasks in animal models (27, 28, 32–35), but on visual discrimination in human tasks, which hinders the proper data transposition from animal models to humans and vice versa (36, 37). For example, it has been demonstrated that the neurobiological processes at work in a reversal learning task differ according to the type of stimuli used with cross-species inconsistencies due to the use of different types of stimuli (38).

Therefore, in order to study the involvement of behavioral flexibility in compulsive behaviors, we propose to assess this cognitive function through a comparative approach in both OCD patients and the *Sapap3* KO mouse model. For this purpose, we have developed an innovative high throughput behavioral setup for mice that allows us to reliably test multiple times the performance of individual subjects in a non-spatial visual reversal learning task, as it is commonly performed in human studies. Finally, we ensured the correct interpretation of our results by recruiting large samples of well characterized and selected subjects in both species, thereby enabling us to look at intra-species variability in our analyses.

## METHODS AND MATERIALS

### Participants and animal subjects

Forty patients diagnosed with OCD according to the DSM-V criteria and with a score greater than or equal to 16 on the Yale-Brown Obsessive-Compulsive Scale (YBOCS) were included in this study (see Supplement for details). Additionally, the current pharmacological treatment was converted to dose-equivalent fluoxetine for each patient (39). Forty healthy control subjects were matched individually according to age, sex, handedness, school education as well as for IQ (estimated by the French National Adult Reading Test, fNART (40)). The protocol for human participants was approved by the Medical Ethical Review Committee of the Pitié-Salpêtrière Hospital (ID RCB n° 2012-A01460-43).

Fifty-two C57BL/6J male mice (26 Sapap3-null (KO) and 26 age matched wildtype (WT) littermates), 6-7 months old, were used. Each animal experiment was approved by the Ethics committee Darwin/N°05 (Ministère de l’Enseignement Supérieur et de la Recherche, France) and conducted in agreement with institutional guidelines, in compliance with national and European laws and policies (Project n° 00659.01).

The detailed characteristics of the samples for both species are summarized in Table 1 (see Supplement for details).

**Table 1.** Demographic and clinical characteristics of the samples. Mean (SD). A BF_10_ greater than one is in favor of a difference and vice versa. The further the BF_10_ is from one, the greater the evidence and vice versa. The * indicates a JZS ANOVA BF_10_ with the results of post hoc tests given after the BF_10_ value; “=” indicating an absence of difference. AAP: atypical antipsychotic.

| Humans |  |  |  |  |  |  |
| --- | --- | --- | --- | --- | --- | --- |
| | Healthy (H) | OCD | $BF_{10}$ | Checkers (C) | Non-checkers (NC) | $BF_{10}$ |
| Sample size | 40 | 40 |  | 21 | 19 |  |
| Gender (M/F) | 15/25 | 15/25 | 0.26 | 10/11 | 5/14 | 0.44 |
| Hand (R/L) | 35/5 | 34/6 | 0.2 | 16/5 | 18/1 | 0.34 |
| Age in years | 40.28 (13.59) | 40.15 (13.22) | 0.18 | 41.9 (12.78) | 38.21 (13.77) | 0.15* |
| Education in years | 5.43 (2.19) | 5.15 (2.69) | 0.37 | 5.1 (3.24) | 5.21 (1.99) | 0.13* |
| fNART IQ | 108.18 (6.76) | 107.4 (6.87) | 0.2 | 106.81 (7.12) | 108.05 (6.71) | 0.14* |
| BDI | 0.8 (1.34) | 12.53 (6.56) | >100 | 11.05 (5.09) | 14.16 (7.68) | >100*: H < C = NC |
| BIS-10 | 45 (9.89) | 47.03 (11.75) | 0.23 | 48.57 (12.14) | 45.32 (11.38) | 0.21* |
| STAI-A | 40.40 (4.38) | 62.60 (13.07) | >100 | 62.81 (10.08) | 62.37 (16.04) | >100*: H < C = NC |
| STAI-B | 36.05 (5.97) | 65.38 (9.05) | >100 | 64.76 (5.65) | 66.05 (11.88) | >100*: H < C = NC |
| OCI-R: |  |  |  |  |  |  |
| Total score | 3.88 (4.23) | 30.43 (8.86) | >100 | 31.19 (7.85) | 29.58 (10) | >100*: H < C = NC |
| Order subscore | 1.45 (1.88) | 6.28 (3.06) | >100 | 6.14 (3.14) | 6.42 (3.04) | >100*: H < C = NC |
| Washing subscore | 0.53 (1.06) | 6.45 (4.59) | >100 | 4.43 (4.23) | 8.68 (3.96) | >100*: H < C < NC |
| Checking subscore | 0.35 (0.89) | 5.68 (3.87) | >100 | 8.52 (2.23) | 2.53 (2.65) | >100*: H < NC < C |
| Hoarding subscore | 1.1 (1.37) | 3.85 (2.93) | >100 | 4.29 (2.26) | 3.37 (3.53) | >100*: H < C = NC |
| Obsession subscore | 0.38 (0.77) | 5.53 (2.97) | >100 | 5.1 (2.41) | 6 (3.5) | >100*: H < C = NC |
| Neutralization subscore | 0.18 (0.5) | 2.65 (2.86) | >100 | 2.71 (3.02) | 2.58 (2.76) | >100*: H < C = NC |
| YBOCS |  | 25.25 (5.29) |  | 24 (4.84) | 26.63 (5.54) | 0.85 |
| Taking an antidepressant: |  | 28 |  | 16 | 12 | 0.74 |
| with an AAP |  | 4 |  | 2 | 2 | 0.24 |
| with a benzodiazepine |  | 7 |  | 4 | 3 | 0.3 |
| Fluoxetine-equivalent in mg/d |  | 53.73 (57.65) |  | 46.74 (47.88) | 61.45 (67.34) | 0.4 |
| Psychiatric comorbidities: |  |  |  |  |  |  |
| Anxiety disorder |  | 11 |  | 9 | 2 | 6.27 |
| Eating disorder |  | 1 |  | 0 | 1 |  |
| Age at onset |  | 15.23 (5.88) |  | 15.05 (5.64) | 15.42 (6.28) | 0.31 |
| Duration in years |  | 24.92 (13.95) |  | 26.86 (15.45) | 22.79 (12.14) | 0.43 |
| C57BL/6J Mice |  |  |  |  |  |  |
| | WT (W) | *Sapap3* KO | $BF_{10}$ | Impaired KO (I) | Unimpaired KO (U) | $BF_{10}$ |
| Sample size | 26 | 26 |  | 12 | 14 |  |
| Age in days | 200.12 (12.23) | 199.35 (12.29) | 0.25 | 199.67 (14.9) | 199.07 (10.13) | 0.16* |
| Weight in g | 35.47 (4.99) | 29.71 (2.88) | >100 | 29.13 (3.32) | 30.21 (2.46) | >100*: W > I = U |
| Number of grooming bouts | 5.38 (3.75) | 12.12 (7.05) | >100 | 14.58 (8.55) | 10 (4.82) | >100*: W < I = U |
| Time spent grooming as % | 11.76 (19.18) | 25 (20.43) | 3.11 | 29.01 (24.2) | 21.56 (16.72) | 1.71*: W < I = U |

### Mouse automatized experimental chamber

It has been shown that daily manipulation of these genetically-modified anxious animals in experimental procedures can increase stress and negatively impacts on behavioral results, including the assessment of behavioral flexibility (41). To avoid this bias, especially in *Sapap3* KO mice which express an anxious phenotype, we designed and used in our study an automated experimental chamber (Figure 1A) where mice were exposed to the task 24h a day. The mouse lived in the experimental chamber for several weeks where water was provided *ad libitum*, and each trial could be self-initiated to get food. Therefore, these in-house experimental chambers allow a more ecological assessment of behavioral performance by avoiding any daily manipulation and methodological constraints (such as food deprivation) that could affect animals normal behavior (42–44). Moreover, our device allowed, as in humans, to repeat the measurement of interest through the collection of thousands of trials per mouse (Figure 1B; see Supplement for details). To control for any environmental influence, the mice underwent the task in pairs, a KO and its WT littermate starting at the same time. Prior to the beginning of the task itself, the mice were acclimatized to the experimental chamber for 24 hours with a pellet delivered each time they poked their nose into the food hopper. After this acclimatization phase, the master program automatically triggered two successive pre-training instrumental phases where the animal learned to activate a display and to respond to visual stimuli. Finally, when the animal reached pre-defined criteria of completion of these pre-training phases, the reversal learning task could proceed (Supplementary Figure S1).

**Figure 1.**
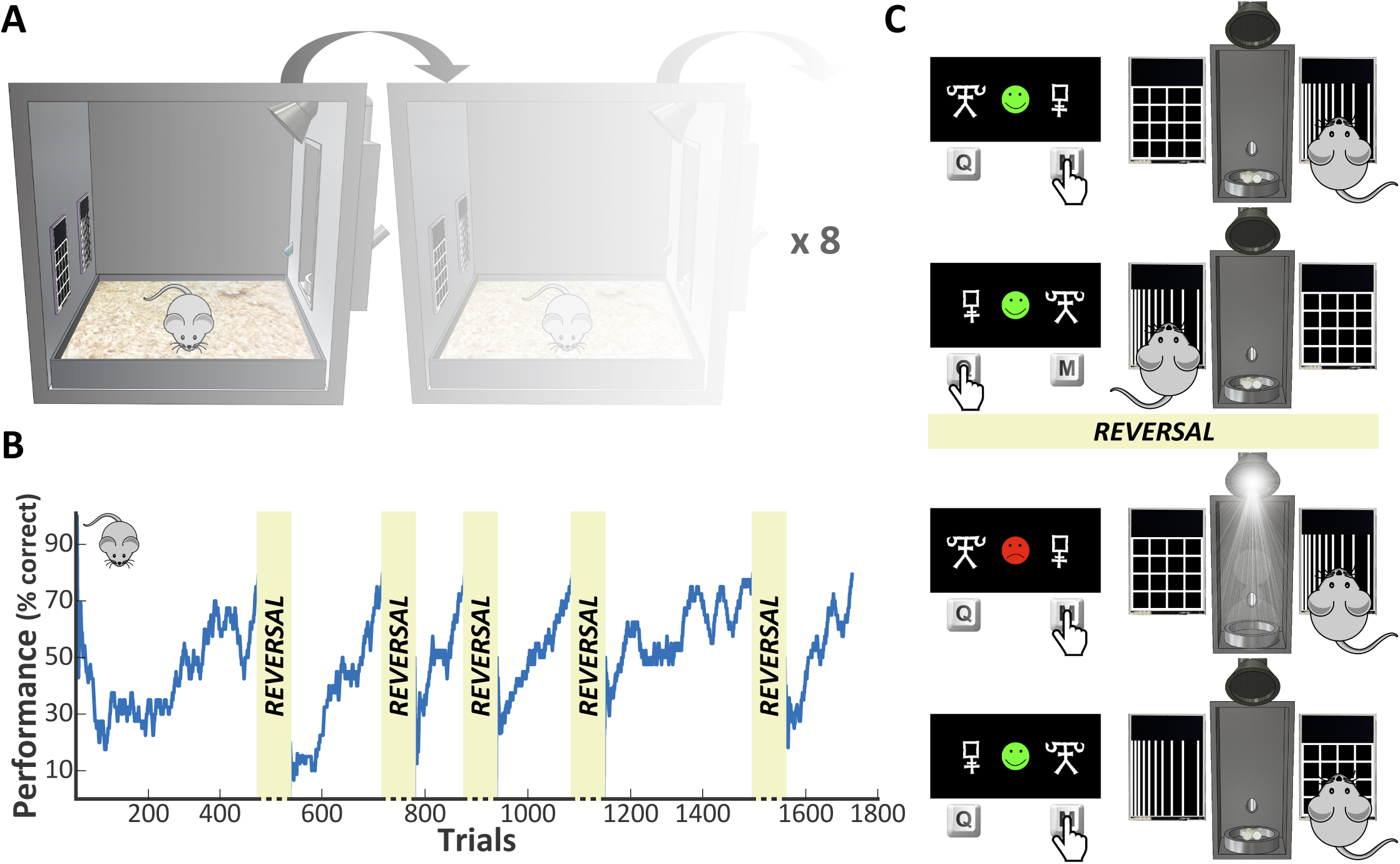
The reversal learning task. **(A)** Illustration of the behavioral apparatus. Up to 8 operant conditioning chambers run in parallel with mice living and working without any human intervention. Left of each box: capacitive touch-sensitive screens. Front-right of each box: the pellet dispenser. Rear-right of each box: the water dispenser. Upper right of each box: LED illumination. **(B)** Example of the performance of one mouse across five reversals. The drop in performance after each reversal indicates that the mouse has correctly learned the previous stimulus-reward association. Blue line: performance within a 40-trial sliding window. **(C)** Design of the human (left) and the mouse (right) versions of the reversal learning task. On each trial, the subject has to make a choice between two different stimuli displayed on the screens. Depending on their choice, positive (for correct response) or negative (for incorrect response) feedback is provided. When the subject has learned the correct association (see detailed criteria in Supplement), the reward contingencies are reversed without warning that the previously positive stimulus become negative (and vice versa).

### Reversal learning paradigm

The human version of the reversal learning task (Figure 1C left) was administered in a computerized version adapted from Valerius et al. (45) and coded in MatLab R2013b (MathWorks) and the Psychophysics Toolbox v3 module. As classically done in human behavioral studies (46), to make reversal events less obvious, probabilistic errors were interspersed so that there was a 20% chance of receiving misleading feedback. The task ended after the completion of 20 reversals. The mouse version of the task (Figure 1C right) was the same as the human version, with the feedback being deterministic as the main difference (i.e. there were no probabilistic error). As in the human task, the rewarded side was contingent to the stimuli presented on the screens to exclude any simple spatially-guided or even cue-guided strategy. The task ended after completion of 5 reversals, representing around 2000 of trials performed over approximately 3 weeks (Figure 1B and Supplementary Figure S2).

For both species, the main behavioral measures to assess the subjects’ performance were the number of trials needed to reach the reversal criterion and the number of perseverative errors following a reversal (reversal errors). For mice, this last measure was estimated as a proportion of errors in the perseverative phase (47, 48) which was defined as a block of trials above a 60% error rate following a reversal event. These perseverative trials were determined according to a change point analysis adapted from Gallistel et al. (49). Other parameters of interest were the probability of a spontaneous strategy change (SSC, i.e. switching to the alternative unrewarded stimulus despite positive feedback), also defined as response lability; and the number of perseverative errors following a SSC (SSC errors) (see Supplement for details).

### Statistical analysis

Bayesian statistics were used to overcome the multiple shortcomings of *null hypothesis significance testing* (50). This approach allows us to assess not only the strength of the evidence against the null hypothesis but also the one in favor of it by computing the Bayes Factor (51), a ratio that contrasts, given the data, the likelihood of the alternative hypothesis (H_1_) with the likelihood of the null hypothesis (H_0_), hence noted BF_10_. BF values have a natural and straightforward interpretation as indicative of “substantial” (3<BF<10), “strong” (10<BF<30), “very strong” (30<BF<100) and “decisive” (BF>100) evidence in favor of H_1_ (and conversely for H_0_ for BF values below ^1^/_3_, ^1^/_10_, ^1^/_30_ and ^1^/_100_ respectively). All the analyses were performed in JASP v0.9.2 (52). For group comparisons, two-tailed (or one-tailed when justified, indicated by BF_±0_) Jeffreys–Zellner–Siow (JZS) paired t-tests (53) were carried out to analyze differences in continuous variables with an uninformed Cauchy prior (µ = 0, σ = ^1^/_√2_), and two-tailed Gunel-Dickey (GD) contingency tables tests (54) for independent multinomial sampling were carried out to analyze differences in categorical variables with a default prior concentration of 1. For multiple-group comparisons, we used JZS ANOVA (55) with an uninformed multivariate Cauchy prior (µ = 0, σ = ^1^/_2_) followed by post-hoc two or one-tailed JZS t-tests and two-tailed GD contingency tables tests for joint multinomial sampling. When appropriate, effect sizes were reported, i.e. Cohen’s *d* with its 95% Credible Interval (95CI) for JZS t-tests and the η^2^ for JZS ANOVA.

To assess the inter-dependence between compulsive behavior severity and behavioral flexibility performance, we carried out correlation analyses between the severity score of the disorder and the behavioral parameters extracted from our task, as performed in previous studies in both species (35, 45). In humans, as the different OCD clinical subtypes can have a distinct impact on behavioral flexibility (19), we analyzed separately the relationship between these subtypes, measured by the OCI-R subscores, and behavioral flexibility. Likewise, we additionally explored the influence of depressive and anxious symptoms as well as antidepressant dose on task parameters. These analyses relied on two-tailed JZS Pearson correlation tests (56) with an uninformed stretched β prior (width 1). The correlation coefficient *r* is reported along with its 95CI. The categorical variables are expressed as percentages and the continuous variables are expressed as means ± standard deviation. All values are rounded to two decimal places.

### Cluster analysis

To search for a potential sub-group of WT or KO mice which might behave differently in our task, we performed a two-step cluster analysis (57) using the four behavioral parameters extracted from our task (number of trials to reversal, reversal errors, SSC probability and SSC errors). This algorithm has the advantage of relying on the Bayes Information Criterion (BIC) to automatically choose the number of clusters to retain; avoiding any subjective and biased selection. Two-step clustering also offers an overall goodness-of-fit measure called the silhouette measure of cohesion and separation. A silhouette measure of less than 0.20 indicates a poor solution quality, a measure between 0.20 and 0.50 a fair solution, whereas values of more than 0.50 indicate a good solution. Furthermore, the procedure indicates each variable’s importance for the construction of a specific cluster with a value ranging from 0 (least important) to 1 (most important). Our clustering used log-likelihood as a measure of distance and the BIC for automatic determination of the number of clusters. It was followed by a stepwise discriminant analysis as a confirmatory procedure and included all mice (KO and WT). The analysis used the Wilks’ lambda for variable selection and prior probabilities computed from group sizes along with the within-groups covariance matrix for classification. These analyses were performed in SPSS v25 (IBM) (See Supplement for more details).

## RESULTS

### Compulsiveness is unrelated to behavioral flexibility as assessed by the reversal learning paradigm

In both species, the performance profile after a reversal event was similar between the compulsive subjects and their controls (two-way mixed ANOVA, BF_Inclusion_ < 1 for group factor, Figure 2A and Supplementary tables S1 and S2). Considering the number of trials needed to reach the reversal criterion (Figure 2B up), no group differences were found for both humans (BF_10_ = 0.64, *d* = 0.27 [0.04 0.77]) and mice (BF_10_ = 0.7, *d* = 0.33 [-0.1 0.92]). Similarly, no significant group differences were found considering the number of reversal errors (Figure 2B down) for both humans (BF_10_ = 1.5, *d* = 0.35 [0.07 0.88]) and mice (BF_10_ = 0.44, *d* = 0.25 [-0.2 0.83]). The comparison of other behavioral parameters, such as SSC and SSC errors, support this lack of difference between groups for both species (Table 2).

**Table 2.** Behavioral parameters. Mean (SD). A BF_10_ greater than one is in favor of a difference and vice versa. The further the BF_10_ is from one, the greater the evidence and vice versa. For JZS ANOVA, the results of *post hoc* tests are given after the BF_10_ value with “=” indicating an absence of difference.

| Humans |  |  |  |  |  |  |
| --- | --- | --- | --- | --- | --- | --- |
| | Healthy (H) | OCD | $BF_{10}$ | Checkers (C) | Non-checkers (NC) | $BF_{10}^{ANOVA}$ |
| **Trials in acquisition phase** | 19.79 (7.29) | 24.09 (9.89) | 2.29 | 28.13 (11.59) | 19.62 (4.74) | 47.88: C > NC = H |
| **Trials to reversal** | 26.09 (6.84) | 29.09 (8.43) | 0.64 | 31.87 (10.05) | 26.02 (4.8) | 3.98: C > NC = H |
| **Reversal errors** | 1.64 (0.4) | 1.48 (0.27) | 1.5 | 1.46 (0.26) | 1.51 (0.28) | 0.72 |
| **SSC probability as %** | 4.1 (3.17) | 5.2 (4.68) | 0.41 | 6.47 (5.44) | 3.8 (3.27) | 1.19: C > NC = H |
| **SSC errors** | 0.17 (0.19) | 0.17 (0.16) | 0.17 | 0.21 (0.17) | 0.12 (0.15) | 0.35 |
| **SCAPE probability as %** | 54.79 (22.53) | 61.82 (15.26) | 0.6 | 61.86 (14.78) | 61.77 (16.17) | 0.33 |
| **SCAPE errors** | 1.28 (0.16) | 1.29 (0.21) | 0.19 | 1.31 (0.25) | 1.26 (0.15) | 0.16 |
| C57BL/6J Mice |  |  |  |  |  |  |
| | WT (W) | *Sapap3* KO | $BF_{10}$ | Impaired KO (I) | Unimpaired KO (U) | $BF_{10}^{ANOVA}$ |
| **Trials in acquisition phase** | 464.66 (182.23) | 438.39 (248.19) | 0.22 | 560.08 (261.76) | 334.07 (187.31) | 2.35: U < I = W |
| **Trials to reversal** | 437.7 (156.89) | 554.39 (290.37) | 0.7 | 746.87 (278.55) | 389.4 (181.61) | >100: W = U < I |
| **Reversal errors as %** | 25.84 (12.8) | 22.08 (12.97) | 0.44 | 11.1 (5.17) | 31.49 (9.71) | >100: I < U = W |
| **SSC probability as %** | 44.19 (3.65) | 44.4 (4.09) | 0.21 | 48.1 (1.88) | 41.22 (2.36) | >100: U < W < I |
| **SSC errors** | 0.57 (0.11) | 0.45 (0.09) | 27.74 | 0.41 (0.08) | 0.48 (0.1) | >100: I < U < W |

**Figure 2.**
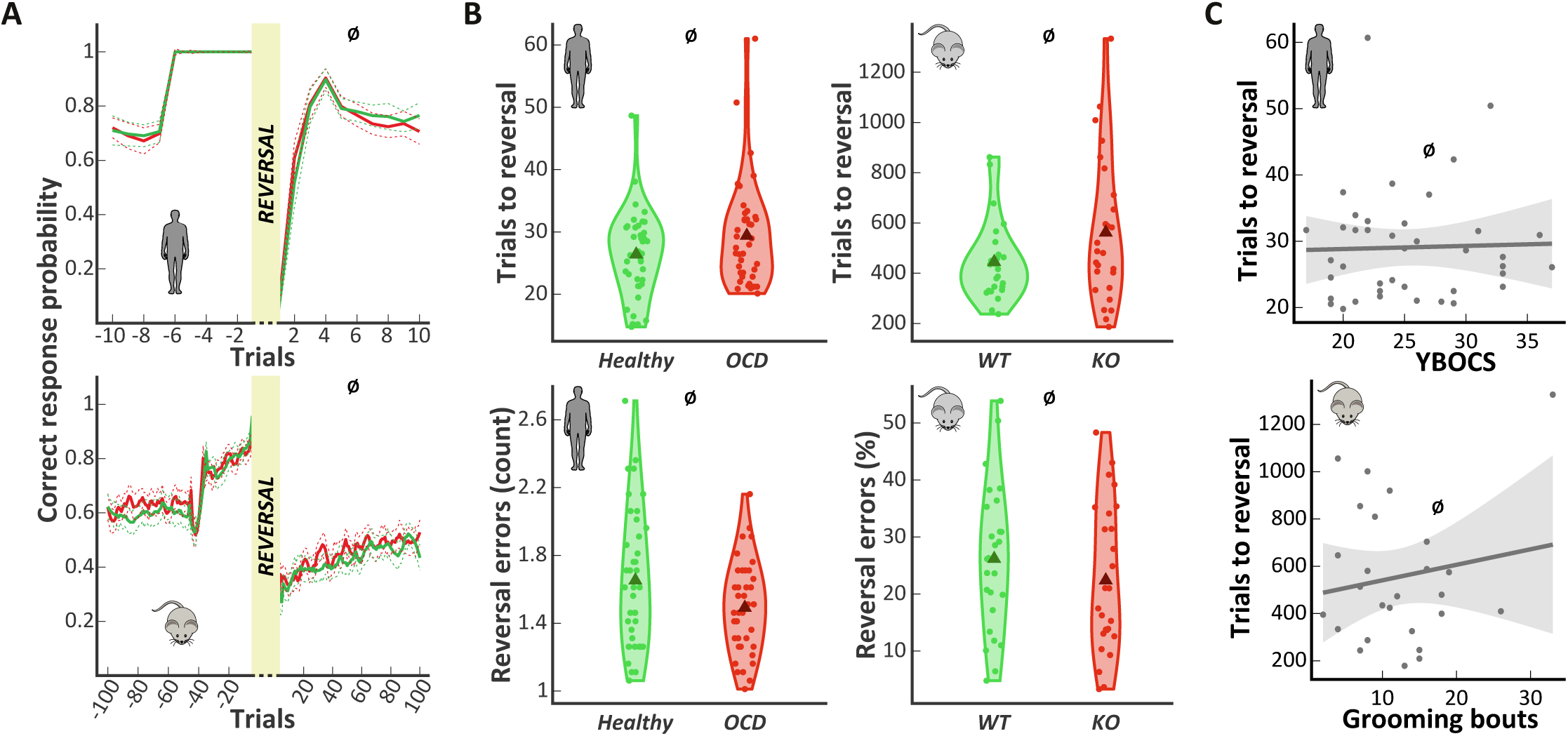
Compulsivity is not related to behavioral flexibility. **(A)** Changes in correct response probability around a reversal. **Above:** for humans with 10 trials around the reversal. *Red line: OCD patients. Green line: healthy subjects. The Savitzky-Golay smoothing algorithm was applied to the data.* **Below:** for mice with 100 trials around the reversal. *Red line: Sapap3* KO mice. *Green line: WT mice. The Savitzky-Golay smoothing algorithm was applied to the data*. **(B)** No difference was found between groups when taken as a whole for both humans (left, n = 40 per group) and mice (right, n = 26 per group), neither when considering the number of trials needed to reach the reversal criterion (above) or the number of reversal errors (below). *Triangle: group mean. Dot: individual mean.* ^*Ø*^: *BF*_*10*_ *< 1.* **(C) Above:** In OCD patients, the disease severity assessed by the Y-BOCS does not predict the number of trials needed to reach the reversal criterion. *Dark line: linear fit. Gray area: confidence interval.* **Below:** In *Sapap3* KO mice, compulsive grooming severity assessed by the number of grooming bouts initiated in a 10-minute period does not predict the number of trials needed to reach the reversal criterion. *Dark line: linear fit. Gray area: confidence interval.*

In OCD patients, correlation analysis showed that disease severity and task performance were not related (Figure 2C up and table 3). The same observation was made for mice with no correlation found between grooming level and the main behavioral parameters (Figure 2C down and Table 3).

**Table 3.**
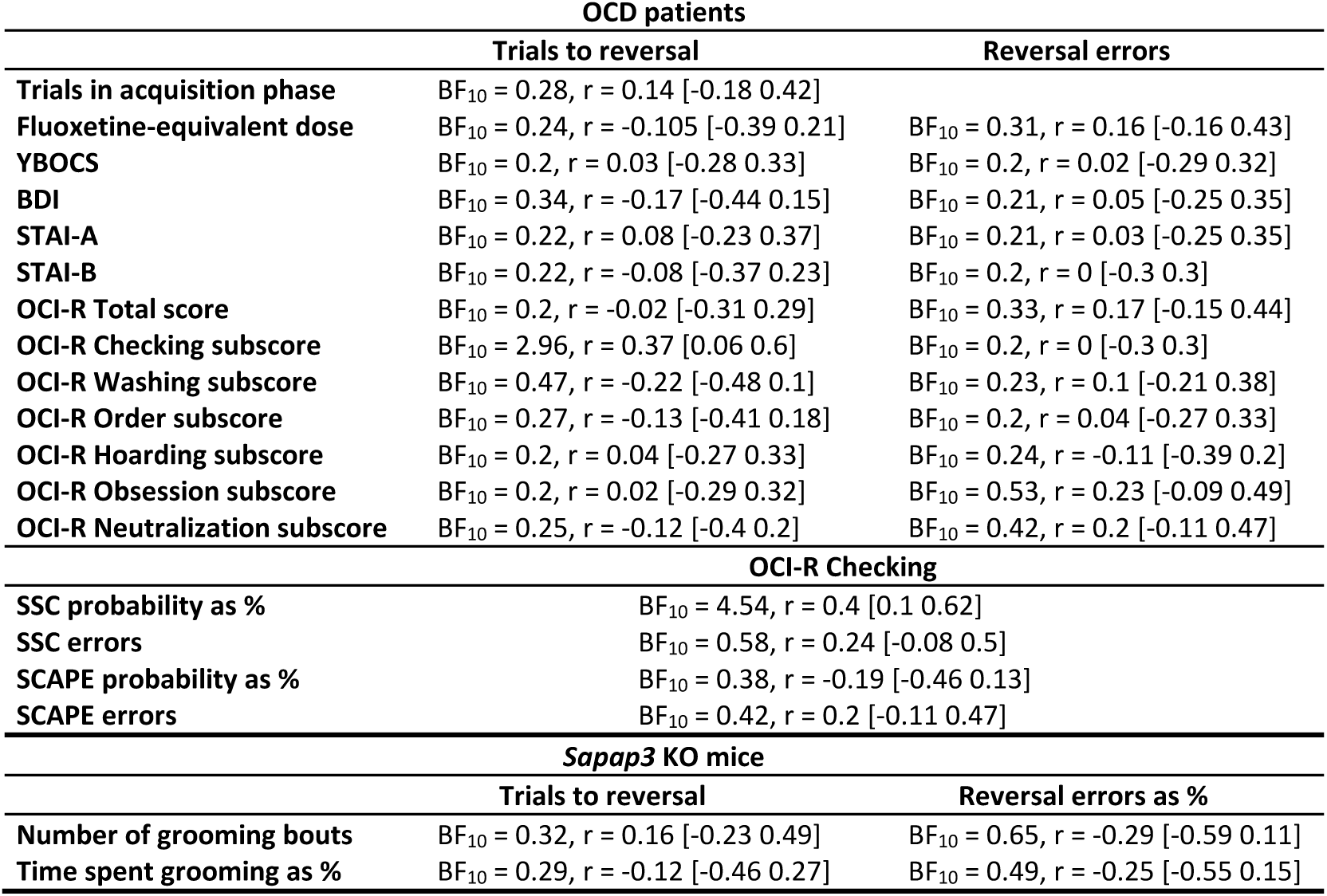
Correlation matrix. A BF_10_ greater than one is in favor of a correlation and vice versa. The further the BF_10_ is from one, the greater the evidence and vice versa.

### Distinct subgroups of OCD patients and KO mice exhibit a behavioral flexibility deficit

In OCD patients, depression/anxiety levels and antidepressant dose does not influence task performance (Table 3). When we assessed the effect of certain symptom subtypes on task performance, only checking symptom severity was related to an increased number of trials needed to reach the reversal criterion (Figure 3A and Table 3). We thus conducted a subgroup analysis by separating the OCD patients with predominantly checking symptoms from the others. Twenty-one OCD “checkers” were separated from the original group. The three resulting new groups (OCD “checkers”, OCD “non-checkers” and healthy subjects) were not different in terms of demographic characteristics (Table 1). In terms of their clinical characteristics, the OCD “checkers” group showed a higher rate of comorbid anxiety disorder compared to the others (Table 1).

**Figure 3.**
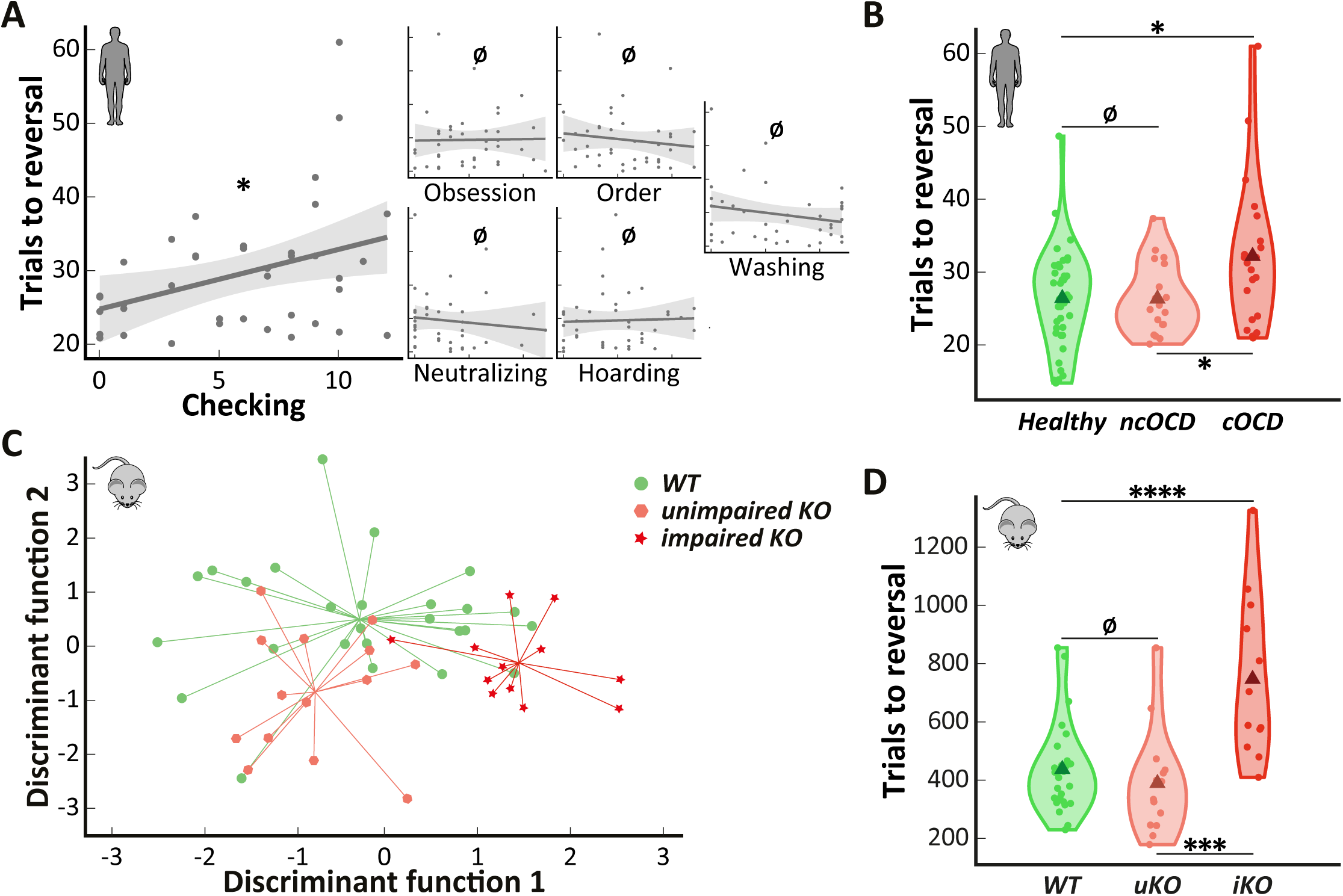
Only a subgroup of OCD patients and KO mice is impaired. **(A)** Only the severity of the checking symptoms measured by the OCI-R checking subscore predicts the number of trials needed to reach the reversal criterion. The higher the checking symptoms severity, the higher the number of trials needed. *Dark line: linear fit. Gray area: confidence interval*. **(B)** Based on this result, we segregated OCD patients with predominant checking symptoms from the others and found that only these were impaired in terms of trials required to reach the reversal criterion. *ncOCD: “non-checkers”. cOCD: “checkers”. Triangle: group mean. Dot: individual mean.* **(C)** A stepwise discriminant analysis was performed on the whole mouse sample on the basis of the four behavioral parameters extracted from our task (number of trials to reversal, reversal errors, SSC probability and SSC errors) to confirm the two clusters identified within the KO mice by the two-step cluster analysis. The classification showed that overall 71.2% of mice were correctly labeled with 83.3% of the “impaired” KO mice correctly classified (16.7% were classified as WTs) vs 50% for the “unimpaired” KO mice (50% classified as WTs). *The intersection point of the lines indicates the group’s centroid.* **(D)** Only the “impaired” KO mice needed more trials to reach the reversal criterion compared to the other KO and WT mice. *uKO: “unimpaired” KO mice. iKO: “impaired” KO mice. Triangle: group mean. Dot: individual mean.* ^*Ø*^*: BF*_*10*_ *< 1. *: BF*_*10*_ *≥ 3. **: BF*_*10*_ *≥ 10. ***: BF*_*10*_ *≥ 30. ****: BF*_*10*_ *≥ 100.*

As for OCD patients, we attempted to determine a similar heterogeneity in the KO mice. Indeed, some innovative studies have found evidence for inter-individual variability of cognitive traits in animal models (58, 59), pointing out it is a necessary consideration for animal studies. To do so, we performed a two-step cluster analysis which identified two clusters within the KO mice with a silhouette measure of 0.6, indicating a good solution. The same analysis was applied to WT controls but no cluster was found. The SSC probability was the most important variable for clusters identification (importance value of 1), followed by the reversal errors proportion (0.79), the number of trials needed to reach reversal criterion (0.42) and the SSC perseverative errors (0.19). Fourteen KO mice were similar to WT in terms of performance and constituted the “unimpaired” group and twelve the “impaired” group. The two KO subgroups were similar in terms of weight at the beginning of the task and grooming level (Table 1), showed similar activity and task engagement (Supplementary Figure S3), and had no identified genealogical difference (Supplementary Figure S4). This clustering was confirmed by the stepwise discriminant analysis: overall 71.2% of mice were correctly labeled with 83.3% of the “impaired” KO mice correctly classified (16.7% were classified as WT) and 50% for the “unimpaired” KO mice (50% classified as WT). These results were consistent with those of the two-step cluster analysis since the “unimpaired” KO mice cluster was closer to the WT mice cluster (Figure 3C).

Considering the number of trials needed to reach the reversal criterion (Table 2), a group effect was found for both humans (BF_10_ = 3.98, η^2^ = 0.11, Figure 3B) and mice (BF_10_ > 100, η^2^ = 0.35, Figure 3D), with an absence of comorbid anxiety disorder effect in humans (Supplementary Table S3). Indeed, OCD “checkers” needed more trials than both OCD “non-checkers” (BF_+0_ = 4.66, *d* = 0.74 [0.08 1.25]) and healthy controls (BF_+0_ = 9.32, *d* = 0.67 [0.14 1.17]); with no difference between the latter two groups (BF_10_ = 0.28, *d* = 0.01 [-0.5 0.5]). Similarly, “impaired” KO mice needed more trials than WTs (BF_10_ > 100, *d* = 1.37 [0.54 2.17]); with no difference between “unimpaired” KO mice and WTs (BF_10_ = 0.43, *d* = 0.29 [-0.35 0.82]).

### Reversal learning deficit is explained by higher response lability rather than perseveration

Considering that only checking symptoms were associated with a reversal learning impairment in OCD patients, we wanted to see if it could be explained by a greater perseveration. We did not find a correlation between checking symptoms severity and the number of reversal errors (Table 3). This was confirmed by the subgroup analysis with an absence of a group effect on the number of reversal errors (BF_10_ = 0.72, η^2^ = 0.06, Figure 4A left). In mice, we found a group effect relative to the proportion of reversal errors (BF_10_ > 100, η^2^ = 0.34, Figure 4A right) but with “impaired” KO mice making fewer reversal errors than WTs (BF_10_ = 53.84, *d* = 1.51 [0.35 1.93]) and no difference between “unimpaired” KO mice and WTs (BF_10_ = 0.71, *d* = −0.48 [-1.02 0.19]).

**Figure 4.**
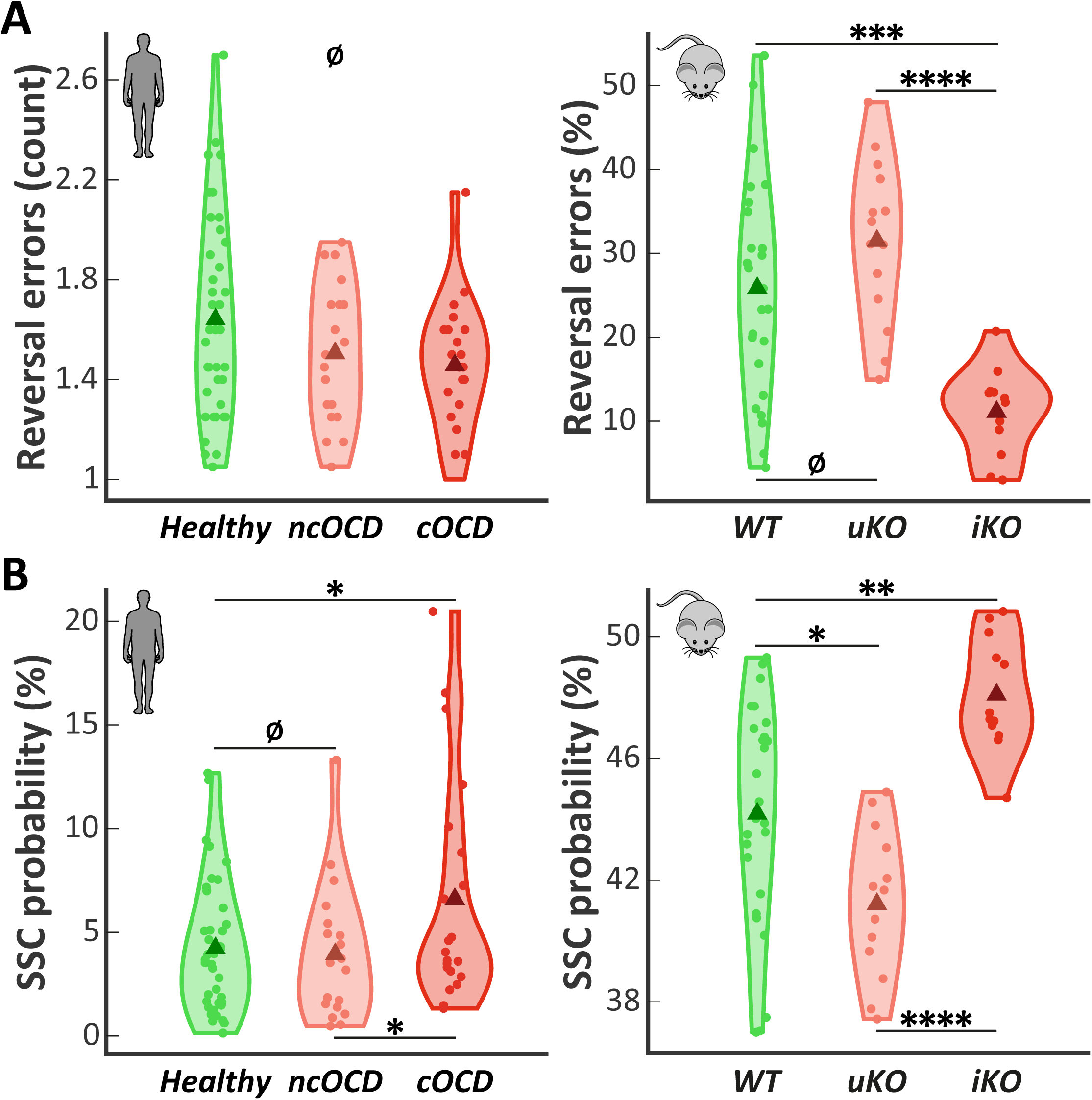
An excessive response lability underlies the reversal learning deficit. The impairment found in both OCD patients and KO mouse subgroups is not explained by **(A)** a greater perseveration, with no difference between the human groups in term of reversal errors and with the “impaired” KO mice making fewer reversal errors than the other KO and WT mice. In fact, it turns out that **(B)** a higher response lability underlies this impairment with both OCD “checkers” and “impaired” KO mice showing an increased SSC probability compared to other groups. *ncOCD: “non-checkers”. cOCD: “checkers”. uKO: “unimpaired” KO mice. iKO: “impaired” KO mice. Triangle: group mean. Dot: individual mean.* ^*Ø*^*: BF*_*10*_ *< 1. *: BF*_*10*_ *≥ 3. **: BF*_*10*_ *≥ 10. ***: BF*_*10*_ *≥ 30. ****: BF*_*10*_ *≥ 100.*

On the contrary, we observed that checking symptoms were associated with a higher response lability in OCD patients with a positive correlation between checking symptoms severity and spontaneous strategy change probability, while no correlation was found with the other parameters (Table 3). The subgroup analysis supported this result with differences between human groups found as we considered the SSC probability (BF_10_ = 1.19, η^2^ = 0.08, Figure 4B left); as for mice (BF_10_ > 100, η^2^ = 0.41, Figure 4B right). OCD “checkers” were more labile than both healthy controls (BF_+0_ = 3.51, *d* = 0.53 [0.08 1.02]) and OCD “non-checkers” (BF_+0_ = 2.22, *d* = 0.59 [0.06 1.1]). The healthy controls and OCD “non-checker” groups did not differ from each other (BF_10_ = 0.29, *d* = 0.09 [-0.41 0.57]) and no comorbid anxiety disorder effect was found (Supplementary Table S4). The same results were obtained in mice with “impaired” KO mice more labile than WTs (BF_10_ = 24.64, *d* = 1.22 [0.33 1.82]), and “unimpaired” KO mice less labile than WTs (BF_10_ = 5.24, *d* = 0.91 [0.12 1.45]). In the same line, WTs perseverated more after SSC, with more SSC errors than “impaired” KO mice (BF_10_ > 100, *d* = 1.61 [0.62 2.24]) (Table 2). Importantly, both OCD “checkers” and “impaired” KO mice show response lability difference only in a reversal context and not during the acquisition phase (see Supplement).

## DISCUSSION

This cross-species study assessed the role of behavioral flexibility in compulsive behaviors through a reversal learning task conducted in both humans and mice. We showed that in both species compulsive subjects do not form a homogeneous group. Taken as a whole, neither the human nor rodent compulsive groups showed differences in task performance compared to their controls. Thus, the severity of compulsive behavior *per se* was not a predictor of performance in our reversal learning task. In contrast, when heterogeneity within groups was taken into account, we found in both species that we could isolate one subgroup showing strong behavioral flexibility deficit in our task, independently of their compulsive behavior severity. In fact, the isolated subgroup of OCD patients was predominantly characterized by checking symptoms. Additionally, we found that this deficit was not underpinned by excessive perseveration behavior after reversal but rather by a greater response lability, again both in humans and mice. Taken together, our results showed from a cross-species perspective that a link between compulsion and behavioral flexibility is not supported but rather they suggest that other dimensions found in subgroups of compulsive subjects, such as excessive response lability, have an effect on behavioral flexibility. Our study also emphasizes the importance of considering clinical subtypes within OCD patients, as encouraged by other recent studies (60, 61).

The fact that only patients suffering from the checking OCD subtype were impaired in our task is in line with the results of a recent meta-analysis demonstrating that the cognitive profile of checking patients is disrupted more than for other OCD patients (62). Considering the impaired subgroup of *Sapap3* KO mice, this result echoes the one of the recent study of Manning et al. (28) which found that only a subgroup of these mice was impaired in a spatial reversal learning task although no difference in grooming level was highlighted. However, they also found a reduced level of locomotor activity in this subgroup with progressive disengagement from their task. It could be, at least partly, a consequence of the higher anxiety level of the *Sapap3* KO mice (23) induced by daily manipulations which has been shown to skew the behavioral outcomes (41, 44). Despite similar observation concerning reversal learning deficit in a subgroup of *Sapap3* KO in our study, we did not observe such a decrease of activity of these mice in our “ecological” design. It suggests that the behavioral flexibility deficit repeatedly observed in some *Sapap3* KO mice across studies (27, 28) may originate from processes other than locomotor hypoactivity.

Indeed, we found that for both species this deficit resulted in excessive response lability. This could be seen as a form of perseveration, the subject having difficulties in suppressing the previous association that maintains its influence long after the reversal. However, OCD patients have decision making (63) and information sampling (64) impairments specific to situations of uncertainty. In that respect, this increased response lability could be induced by an increased level of uncertainty provoked by the reversal event. This assumption makes sense when considering the isolated subgroup of “checker” patients displaying a higher degree of uncertainty. To confirm this assumption, it would be interesting to test in the future if uncertainty monitoring and checking behaviors are affected in *Sapap3* KO mice. Indeed, the excessive response lability observed after reversal, when uncertainty is higher, could be interpreted as if these mice compulsively tested to see if the previously rewarded stimulus was still valid.

These cross-species results are a powerful argument in favor of the heterogeneity of cognitive deficits in compulsive disorders and stresses the importance of also considering this heterogeneity in animal models. Indeed, even if inbred mouse lines share identical genetic background, this is not necessarily stable over time and may result in the emergence of new phenotypic traits due to a genetic drift. However, it has been shown that the C57BL/6J strain is one of the strains least susceptible to this effect (65). Furthermore, we could not identify any genealogical specificity for the impaired KO mice subgroup. Another hypothesis for the heterogeneity we observed in our animal model could be of epigenetic and/or environmental origin. It has been shown for example that phenotypic variability can emerge from variations in epigenetic regulation (59, 66–68). Examples include studies on genetically homogeneous WT C57BL/6J mice showing inter-individual variability in the expression of flexible behavior underpinned by variability in serotonin levels within the OFC (69). As we could not identify any subgroup in our WT mice based on the task performance, such inter-individual variability could only be a risk factor whose sole interaction with the *Sapap3* KO mutation leads to an impairment.

In conclusion, we confirm here that OCD is not a homogeneous disorder and therefore the necessity to consider the phenotypic heterogeneity in both patients and animal models to study their cognitive profile. Moreover, we found that compulsive behavior is not necessarily associated with a deficit in behavioral flexibility. In contrast, this study proposes that a behavioral flexibility deficit, only observed in a subset of compulsive subjects, may result from excessive response lability rather than perseveration, both in humans and mice.

## ACKNOWLEDGMENTS AND DISCLOSURES

This work was supported by *Agence Nationale de la Recherche* – ANR-13-SAMA-0013-01_HYPSY (NB, LM, EB); ANR program ERA-NET NEURON JTC 2013_TYMON (LM, EB); *Investissements d’Avenir* program (Labex Biopsy) managed by ANR-11-IDEX-0004-02 (LM, EB) and Program CARNOT Institute/ICM (EB). The authors thank Profs Guoping Feng and Ann Graybiel for generously providing the *Sapap3* KO mice. Dr. Luc Mallet reports grants from *Fondation pour la recherche médicale* (FRM), grants from *Agence Nationale de la Recherche*, grants from *Fondation Privée des HUG*, grants from *Fondation ICM*, grants from Halphen Foundation, outside the submitted work. The other authors report no biomedical financial interests or potential conflicts of interest.

