## Supplementary Material for "A cross-species assessment of behavioral flexibility in compulsive disorders"

### Supplementary Methods and Materials

#### Humans sample

Patients were recruited through an online advertisement posted on a patient association's web site (AFTOC) and among a cohort of severe patients followed in the psychiatric department of Albert Chenevier Hospital. Healthy comparison subjects were recruited through an online advertisement posted on an information web site dedicated to cognitive research (RISC). Diagnoses and co-morbidity were established by an experienced clinician with the French version of the Mini International Neuropsychiatric Interview (MINI v5 (1)). Exclusion criteria were defined as follows: actual major depressive episode, bipolar disorder, acute or chronic psychosis, substance abuse or dependency including alcohol, epilepsy, cerebral injury, or other neurological problems. To assess severity and clinical subtypes of obsessive-compulsive (OC) symptoms, the YBOCS was administered only for OCD patients, and the Obsessive Compulsive Inventory Revised (OCI-R (2)) was used to measure all participants' OC characteristics. Fifteen patients displayed contamination/washing symptoms, thirteen aggressive/checking symptoms, eight predominant aggressive/checking symptoms associated with contamination/washing symptoms and four had predominantly obsessive thoughts, mainly religious/mental rituals. The mean age at onset of OCD symptoms was 15.23 (±5.881) years old and the mean illness duration was 24.92 (±13.951) years. Depression, trait/state anxiety and impulsivity were assessed respectively with the short version of the Beck Depression Inventory (BDI (3)), the Spielberger's State-Trait Anxiety Inventory (STAI (4)) and the Barratt impulsivity scale (BIS-10 (5)) in their French version. Among the patients taking part in this study, twenty-eight were free of any psychiatric comorbidity, eleven had a comorbid anxiety disorder (essentially general and social anxiety disorder) and one an eating disorder. Considering psychotropic medication, twenty-eight patients took an antidepressant drug alone or combined with antipsychotics or mood stabilizer and the remaining patients were medication-free. Control healthy subjects were free of any current psychiatric or neurological disorder and subsequent medications.

#### Mice sample

The mice were born, weaned (at post-natal day 21) and raised in the animal facility of the Brain and Spine Institute. Before performing the task, they were living in group of 3-5 in ventilated cages with ad-libitum access to water and food, under a temperature of 20-22°C and 50-60% humidity, and were maintained under a 12h light/dark cycle (lights on from 8 am to 8 pm). The mouse started the task when they were at least 6 months old to maximize the chance to observe the grooming phenotype without severe skin lesions (6). During the behavioral assessment, which approximately lasted a month, they were single-housed in the experimental cages under the same 12h light/dark cycle until the task ends and with ad-libitum access to water. The first twenty-four hours in the experimental cage consisted in the habituation period. During this period, they had ad-libitum access to food (20 mg precision tablets, 5TUL, Test Diet) and were video-taped from the top in order to quantify the self-grooming behavior (see *Grooming quantification*). After this initial 24h period, mice no longer had access to ad libitum food but a tablet was delivered each time they answered correctly during the behavioral task (see *Behavioral task*). Their weight was monitored every day and they were supplemented with tablets if they weighted under 80% of their initial mass in order to strictly compensate the loss. They were under continuous video-monitoring.

**Grooming quantification**

The recorded videos were manually analyzed using Kinovea v0.8.15. The self-grooming measures (number of grooming bouts and proportion of time spent grooming) were extracted from a ten-minute activity period (a sufficient duration to highlight differences in the self-grooming behavior (7)) started from 8 pm when mice become more active. When a mouse was not active at 8 pm, the time window was moved forward until the mouse wakes up and leaves the nest. When a mouse was already engaged in a grooming behavior at 8 pm, the time window was moved forward to start after of the ongoing grooming sequence. If a mouse was still engaged in a grooming bout at the end of the time window, this one was moved forward in order to include only complete grooming bouts. Self-grooming was defined as one or more of the elements of the syntactic grooming chain in a flexible, non-chained order: elliptical strokes, small strokes, bilateral strokes, flank licks, and tail and genital licks (7–9). Consistent with previous studies, we counted grooming bouts independently when they were separated by more than two seconds (7, 10). Additionally, we considered two grooming bouts as independent when qualitatively different behavior interrupted the grooming sequence (i.e., jumping, locomotion, rearing) (7).

**Humans reversal learning paradigm**

The subjects sat in front of a 17” TFT monitor and a regular keypad. Two different abstract symbols from the Agathodaimon alphabet were displayed in white font on a black background on the left and right visual fields with randomized locations. Subjects had to choose one of these symbols by using either a left (“Q”) or a right (“M”) button-press depending on whether their chosen symbol was on the left or right side of the screen. The symbols remained on screen until the subject made an answer with the instruction to respond as fast as possible. 750 ms after their response, a feedback in the shape of either a green smiling face or a red sad face was displayed during 500 ms indicating whether their answer was correct or not, with the win or loss of one point respectively. Inter-trials intervals were randomly sorted between 750 ms and 1250 ms. After 6 to 15 (randomized) consecutive correct responses, a reversal occurred and subjects had to adapt their response by selecting as correct the formerly wrong symbol. To increase the difficulty of the task, probabilistic errors were interspersed so that there was a 20% chance of receiving a negative feedback despite a correct response and vice versa. In order to avoid misleading continuous probabilistic errors, we set a maximum of 3 possible continuous probabilistic errors and 3 possible probabilistic errors within a 10-trial sliding window. Moreover, probabilistic error never occurred on a reversal event and its following 3 trials. There were 3 breaks of up to 5 minutes during the task with one every 6 reversals. Each break was followed by a change in the displayed pair of symbols. In addition, at random intervals of 3 to 10 trials, subjects were asked to rate their confidence in the answer they just gave, on a scale from one to six. The task ended after the completion of 20 reversals. All participants were first trained on the task with a few trials practice run (the training ended when the reversal criterion was reached) to familiarize themselves with the notion of probabilistic errors.

In addition to the main behavioral parameters (number of trials needed to reach the reversal criterion, reversal errors, SSC probability, SSC errors), two additional parameters compared to mice were also extracted: the probability of a strategy change after a probabilistic error (SCAPE, i.e., switching to the unrewarded stimulus following misleading negative feedback to a correct response) and the number of perseverative errors following a SCAPE (SCAPE errors). The acquisition phase, defined as the trials needed to reach the reversal criterion for the first time, is thought to measure a baseline capacity for learning the associations, and not a reversal learning deficit (11). It was therefore analyzed separately.

**Mice reversal learning paradigm**

Prior to the beginning of the task, the mice were first habituated to the experimental chamber and the food for 24 hours with a pellet delivered each time they nose poked in the pellet receptacle (two deliveries being separated by at least 5 minutes). After this habituation phase, they underwent automatically two phases of instrumental pre-training. In the first phase, the mouse had to learn to touch any of the screen (which were off with no visual stimulus) in order to get a pellet as a reward. The two screens blinked during 15 s after each touch to indicate the availability of a reward. The reward was delivered even if the mouse was not retrieving the reward within this 15 s time period. If the mouse succeeded to do 10 consecutive reward retrievals within 15 s, the second phase was automatically initiated. In the second phase, the mouse had to learn to initiate a trial by doing a nose poke in the pellet receptacle. When they did so, the screens turned on white for 60 s, indicating the mouse to touch one of them. If it did so before the screens turned off, the screen blinked for 15 s as a signal to retrieve the reward. After 10 consecutive successful trials of this second pre-training phase, the reversal learning task was automatically initiated. When the mouse launched a trial by nose poking in the pellet receptacle, two distinct visual pattern of vertical and horizontal bars equally luminescent (see Fig. 1C) were presented in white font on a black background on the left and right touchscreens with pseudo-randomized locations (the same pattern could not appear more than 3 times consecutively or 7 times in a 10-trial sliding window). The equiluminescent stimuli pair was chosen among those recommended by Horner et al. (12), i.e. grid and lines. Once a trial was initiated, the mouse had 60 s to respond before the screens turned off. If the mouse made a correct response, the two screens blinked and the mouse had 15 s to nose poke in the food receptacle for a reward. Otherwise, the aversive light was turned on for 5 s and the stimuli location remained the same for the subsequent trials until a correct response was made (corrective trials to avoid response lateralization). After completing a trial, the mouse could not launch another one within the next 5 s. When the mouse reached a criterion of 80% correct responses over the last 40 trials, a reversal occurred and the mouse had to adapt and switch on choosing the formerly wrong stimulus as the new correct response. The task ended after the completion of 5 reversals. To control for any environmental influence, the mice underwent the task pair by pair, a KO and its WT littermate starting at the same time in different cages. The first rewarding stimulus was counterbalanced between pairs.

**Behavioral apparatus**

We have automatized the behavioral apparatus that the animal could live in the experimental chamber and then be exposed 24h a day to the task without any experimenter intervention. This allowed the mice to work according to its own physiological rhythm without stress and to increase the duration of exposition to the task (between 3 and 4 weeks on average). The behavioral apparatus consisted of a modified ENV-007CTX experimental chamber from Med Associates (Vermont, USA) with interior dimensions of 30.5 x 24.1 x 29.2 cm. The grid floor of the chamber was covered with a stainless-steel tray to receive bedding. On the left wall, there were two 2.8" TFT capacitive touchscreens (#2090, Adafruit, New York, USA) placed laterally and above the bedding tray with each of them controlled by an Arduino (Leonardo model, Adafruit) interfaced with the I/O module (DIG-716B, Med Associates). On the right wall, there was a pellet dispenser (ENV-203-20, Med Associates) delivering 20 mg precision tablets (5TUL, Test Diet, Missouri, USA) into a pellet receptacle (ENV-303WX, Med Associates) equipped with an infrared head entry detector (ENV-303HDW, Med Associates) and centrally placed. On the left of the pellet receptacle, there was a water bottle (ENV-350RMX, Med Associates). On the ceiling, a micro camera (700TVL Super HAD CCD II with a 2.8 mm lens, Sony, Japan) associated with a red LED for night vision (5 mm, 55 cd) was fixed on the center and an aversive light (6W LED spot) was vertically located above the pellet receptacle and tilted toward the touchscreens. The apparatus was controlled by Med-PC IV software (Med Associates) running on a desktop computer (under the Windows 7 OS) equipped with the DIG-700P2-R2 PCI interface card (Med Associates). The task was coded in its proprietary language (MEDState Notation).

**Change point analysis**

To identify the perseverative phase in the mouse version of the task, a change point analysis was performed on the cumulative record of correct responses for each animal and for each reversal block (13). This analysis allowed the detection of change points which marked significant variations in the slope of the cumulative record, a useful metric for identifying changes in performance. We coded in MatLab R2016b (MathWorks) a recursive algorithm based on MatLab functions provided by Gallistel et al. in their 2004 paper (13) to search the individual cumulative records of performance (correct response = 1, incorrect response = 0) for putative change points (the trial deviating maximally from a straight line drawn between the start of the record and the assessed point). A $\chi^2$ test was used to determine whether the frequencies of correct responses between the portion before the putative change point and the portion after it significantly differed. A logit value was derived from it (log of the odds against the null hypothesis that there is no change) to determine the strength of the evidence of a change in performance around the putative change point. A change point was retained if its logit value reached/exceeded the one defined by the user. We ran the algorithm on each reversal block for each animal starting with the highest (and very conservative) logit value of 6 and, as suggested by Rountree-Harrison et al. (14), counting down of 0.1 until we could detect a change point marking a statistically significant distinction between the post reversal perseverative phase which corresponds to a maximum performance level of 40%, and the learning phase which corresponds to a performance level exceeding 40% (15, 16). The algorithm ended when a change point fulfilling the above-mentioned criteria was found or if the logit value reached the lowest acceptable value of 1.3.

**Two-step cluster analysis**

As its name suggests it, this algorithm is based on a two-stage approach: in the first stage, the algorithm undertakes a procedure that is very similar to the k-means algorithm. Based on these results, the procedure conducts a modified hierarchical agglomerative clustering procedure that combines the objects sequentially to form homogenous clusters. This algorithm has the advantage to automatically choose the number of clusters to retain by calculating measures of fit such as the Bayes Information Criterion (BIC).

The silhouette measure of cohesion and separation is essentially based on the average distances between the objects and can vary between -1 and +1.

### Supplementary Results

#### The higher response lability is reversal specific

In order to determine if the increased response lability is reversal specific, we compared the SSC probability in the acquisition phase for both species. In humans, we did not find a group effect ($BF_{10}$ = 0.41, $\eta^2$ = 0.04). In mice, a group effect seemed to exist ($BF_{10}$ = 1.55, $\eta^2$ = 0.12) with no difference between “impaired” KO mice and WT controls (43 ± 4.5% vs 45.54 ± 8.74%, $BF_{10}$ = 0.47, *d* = 0.33 [-0.34 0.88]) and between “impaired” and “unimpaired” KO mice (43 ± 4.5% vs 39.3 ± 6.84%, $BF_{10}$ = 0.92, *d* = -0.63 [-1.24 0.2]) while “unimpaired” KO mice tend to be less labile than WT controls ($BF_{10}$ = 2.44, *d* = 0.77 [0.03 1.31]). Thus, both OCD “checkers” and “impaired” KO mice seem to show response lability only in a reversal context.

### Supplementary Discussion

Although more complex tasks were developed to study cognitive flexibility to a higher degree in humans (17), we chose a reversal learning task in order to measure the same construct in both species. Indeed, the various components of cognitive flexibility are based on various regions of the prefrontal cortex (PFC) (18). However, there are strong divergences between a human and a rodent PFC (19–22). Thus, this task was of particular interest for its ability to measure the simplest and most conserved form of cognitive flexibility through species, namely the ability to reverse a stimulus-reward association (23). Moreover, it is essentially supported for both humans and mice by the OFC (24–26) which is a PFC region shared by both species (27–30), the activity of which is altered in OCD patients whether at rest (31) or during the execution of this task (32). We have also strengthened the cross-species validity of our task by excluding the paradigms classically used in rodents based on spatial discrimination. We rather developed a reversal learning task in mice based on visual discrimination, the most commonly used sensory modality in human studies. Some teams already tried to transpose humans' experimental paradigms to rodents using the same modalities (33–35). However, they remain a minority with scarcely any studies assessing patients and animal models in parallel (36).

Since sight is a less important sense in rodents, some might argue that it would have been wiser to rely on a sensory modality naturally more developed in this species such as smell, to ensure that the impairment is not due to a difficulty in discriminating stimuli. However, beyond the fact that we used visual stimuli validated in rodents (12), the use of species-specific capabilities (as the use of olfactory stimuli) does not allow the generation of data that can be readily generalized to other species (37). On the contrary, the further away from species-specific abilities/behaviors, the more likely it is to assess a function/mechanism that transcends the species barrier (38). Another concern may rely on the use of deterministic feedbacks in rodents rather than probabilistic feedbacks. The choice of probabilistic feedbacks in humans was justified to increase the difficulty of the task as the use of deterministic feedback made it so simple that participants immediately detect the change in contingencies and start responding to the other stimulus. This issue does not arise in rodents for which the task is difficult enough in its deterministic version as it requires hundreds of trials to achieve only one reversal. It has also been shown that this difference is insignificant given the similarity of the data acquired in the two species using this paradigm (39).

In parallel to the effective implementation of a translational approach, one of the strengths of our study lies in the development of a setup dedicated to the behavioral assessment in ecological condition of our mice with a high throughput data acquisition allowing to obtain results unbiased by the stress induced by the environment or the experimenter (40–42). Indeed, most of animal studies relies on food deprivation, multiple labor-intensive sessions with daily manipulations without any respect towards the animal physiological cycle causing stress in animals. These factors are not fully considered with very few teams taking an interest in them and developing fully automated procedures that allow the animal to live and work in the experimental apparatus without these stress factors (43–45).

The use of Bayesian statistics is another strength of our study which allowed us to support the absence of certain differences, notably in terms of "classical" perseveration but also to quantify the weight of evidence in favor or against a difference. This strength also points out a limitation to the interpretation of our results, especially those obtained with humans. Indeed, the differences highlighted between the checking patients and the healthy subjects are all supported by a Bayes Factor lower than 10 reflecting a substantial evidence but far from being decisive with a moderate effect size which is not the case with mice. This points out to the need to replicate these results on an even larger sample than the one included in this study.

### Supplementary tables

**Model Comparison**

| Models | $BF_{10}$ |
|---|---|
| Null model | 1.000 |
| Trial | 3.251e +244 |
| Trial + OCD | 3.651e +243 |
| Trial + OCD + Trial ✻ OCD | 2.683e +243 |
| OCD | 0.094 |

**Analysis of Effects**

| Effects | $BF_{Inclusion}$ |
|---|---|
| Trial | ∞ |
| OCD | 0.130 |
| Trial ✻ OCD | 0.297 |

*Note.* Compares models that contain the effect to equivalent models stripped of the effect. Higher-order interactions are excluded.

**Supplementary Table S1.** OCD did not influence the performance following a reversal event.

**Model Comparison**

| Models | $BF_{10}$ |
|---|---|
| Null model | 1.000 |
| Trial | 4.616e +35 |
| Trial + KO | 1.028e +35 |
| Trial + KO + Trial ✻ KO | 4.600e +30 |
| KO | 0.215 |

**Analysis of Effects**

| Effects | $BF_{Inclusion}$ |
|---|---|
| Trial | ∞ |
| KO | 0.149 |
| Trial ✻ KO | 3.260e -5 |

*Note.* Compares models that contain the effect to equivalent models stripped of the effect. Higher-order interactions are excluded.

**Supplementary Table S2.** The knockout of the *Sapap3* gene did not influence the performance following a reversal event.

**Model Comparison**

| Models | $BF_{10}$ |
|---|---|
| Null model | 1.000 |
| Checking subtype | 3.975 |
| Comorbid anxiety disorder | 0.332 |
| Checking subtype + Comorbid anxiety disorder | 2.069 |
| Checking subtype + Comorbid anxiety disorder + Checking subtype ✻ Comorbid anxiety disorder | 1.250 |

**Analysis of Effects**

| Effects | $BF_{Inclusion}$ |
|---|---|
| Checking subtype | 4.539 |
| Comorbid anxiety disorder | 0.483 |
| Checking subtype ✻ Comorbid anxiety disorder | 0.604 |

*Note.* Compares models that contain the effect to equivalent models stripped of the effect. Higher-order interactions are excluded.

**Supplementary Table S3.** Effect of having a comorbid anxiety disorder on the mean number of trials needed to reach reversal criterion.

**Model Comparison**

| Models | $BF_{10}$ |
|---|---|
| Null model | 1.000 |
| Checking subtype | 1.189 |
| Comorbid anxiety disorder | 0.361 |
| Checking subtype + Comorbid anxiety disorder | 0.444 |
| Checking subtype + Comorbid anxiety disorder + Checking subtype ✻ Comorbid anxiety disorder | 0.163 |

**Analysis of Effects**

| Effects | $BF_{Inclusion}$ |
|---|---|
| Checking subtype | 1.200 |
| Comorbid anxiety disorder | 0.367 |
| Checking subtype ✻ Comorbid anxiety disorder | 0.367 |

*Note.* Compares models that contain the effect to equivalent models stripped of the effect. Higher-order interactions are excluded.

**Supplementary Table S4.** Effect of having a comorbid anxiety disorder on the probability of spontaneous strategy change.

### Supplementary figures

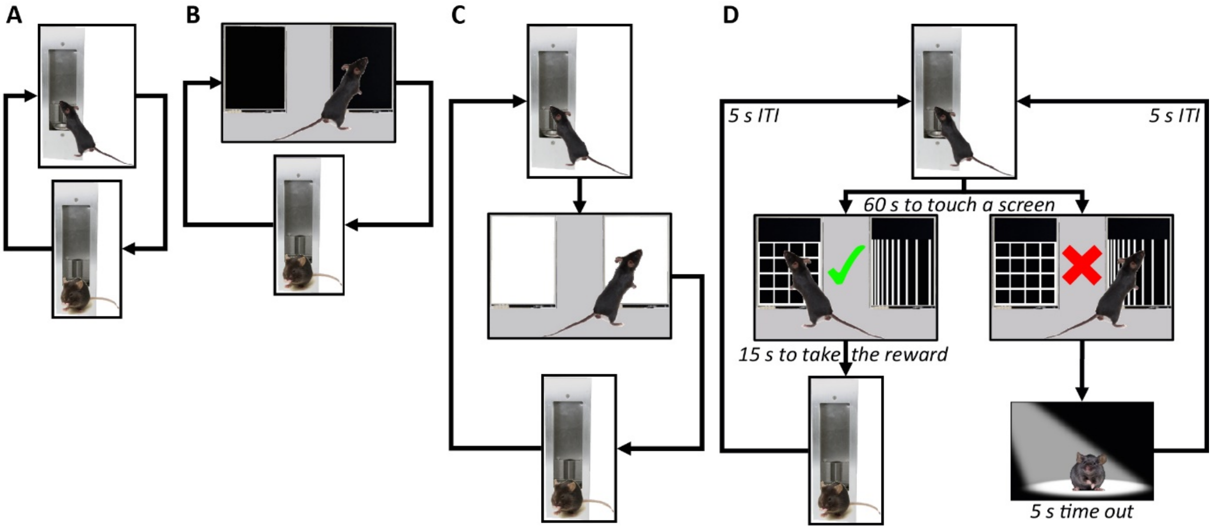


**Supplementary Figure S1.** The four stages of the mouse version of the task. **(A)** Habituation to the experimental chamber and the food for 24 hours. *A pellet is delivered each time the mouse nose poke in the pellet receptacle (two deliveries being separated by at least 5 minutes).* **(B)** Screen touch learning: the mouse has to learn to touch the screens in order to get a pellet as a reward. *One screen touch initiates a 15-second blink to indicate the reward availability. This stage ends with 10 rewards consecutively retrieved within 15 seconds.* **(C)** Trial initiation learning: the mouse has to learn to launch a trial before touching a screen in order to get a reward. *A nose poke in the pellet receptacle turns on white the screens for 60 s, indicating the mouse to touch one of them. A screen touch within 60 s turns them off and launch a 15-second blink to indicate the reward availability. This stage ends with 10 rewards consecutively retrieved within 15 s.* **(D)** A trial sequence of the reversal stage. *First, the mouse has to nose poke into the pellet receptacle to launch a trial, triggering the stimuli display. It has then 60 s to choose a stimulus, otherwise the screens turn off and the mouse will have to launch a trial again. In case of a correct response, the mouse has 15 s to retrieve the pellet. Otherwise, the aversive light is turned on for 5 seconds. After completing a trial, the mouse could not launch another one within the next 5 s.*

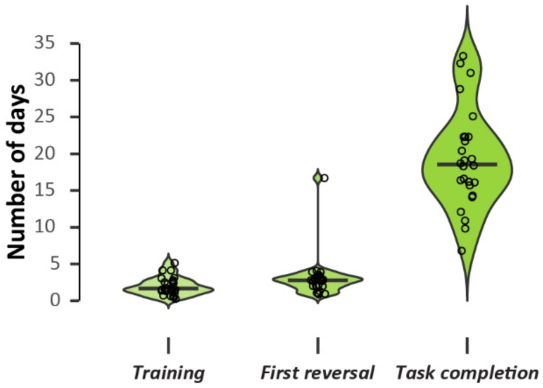


**Supplementary Figure S2.** Average number of days required to complete the different phases of the task. *n = 26 WT. Bar: median. Circles: individual data.*

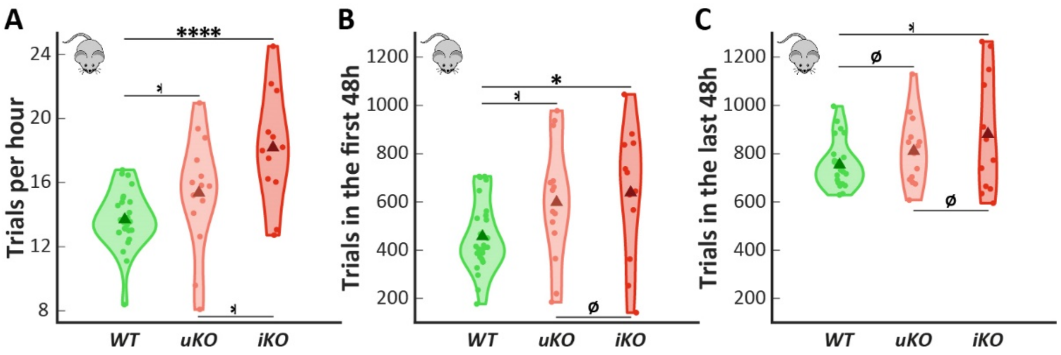


**Supplementary Figure S3.** The *Sapap3* KO mice are more active compared to WT mice. **(A)** The *Sapap3* KO mice performed more trials per hour in average than WT mice with the impaired KO mice subgroup having the highest rate. **(B)** The *Sapap3* KO mice performed more trials in the first 48 hours after the beginning of the task. **(C)** No significant difference in engagement level in the last 48 hours of the task (3 weeks after the beginning of the task in average). All mice were significantly more engaged in the last 48 hours compared to the first 48 hours (JZS two-way mixed ANOVA, $BF_{10} > 100$, $\eta^2 = 0.5$; with no group influence over time, $BF_{10} = 0.99$, $\eta^2 = 0.02$). *uKO: "unimpaired" KO mice. iKO: "impaired" KO mice. Triangle: group mean. Dot: individual mean.*

*ø: $BF_{10} < 1$. ⁎: $BF_{10} > 1$. *: $BF_{10} \geq 3$. **: $BF_{10} \geq 10$. ***: $BF_{10} \geq 30$. ****: $BF_{10} \geq 100$.*

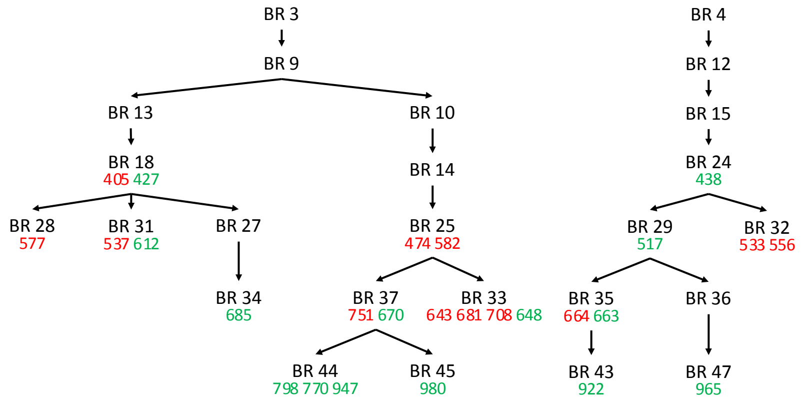


**Supplementary Figure S4.** *Sapap3* KO mice genealogy. *BR = breeding pair. In green: unimpaired KO mouse. In red: impaired KO mouse.*
